# The Orsay Virus as a model for population-wide viral infection dynamics

**DOI:** 10.1101/2021.03.02.433572

**Authors:** Laurence Pirenne, Maximilian A. H. Jakobs, David Jordan, Kristian Franze, Eric A. Miska

**Author notes:** Corresponding author: Eric A. Miska., **Email:**. Authors contributed equally (Laurence Pirenne, Maximilian A. H. Jakobs). **Classification** Biological Sciences, Genetics. Author Contributions LP, DJ, and MAHJ performed experiments MAHJ, LP, and DJ analyzed data., LP and MAHJ wrote the original manuscript draft All authors reviewed and edited the manuscript. EM and DJ conceptualized the project, EM and KF administrated and supervised the project.

## Abstract

To this day, epidemics pose a considerable threat to mankind. Experimental models that simulate the spread of infectious diseases are thus crucial to the inception of effective control policies. Current models have had great success incorporating virulence and host immune response but do rarely take host genetics, behavior and host environment into account. Here, we present a full-scale imaging setup that utilizes the infection of the nematode *C. elegans* with a positive-stranded RNA virus (Orsay Virus) to probe key epidemiological parameters and simulate the spread of infection in a whole population. We demonstrate that our system is able to quantify infection levels and host behavior at a high sampling rate and show that different host genetic backgrounds can influence viral spread, while also highlighting the influence of infection on various host behaviors. Future work will allow the isolation of key behavioral and environmental factors that affect viral spread, potentially enabling novel policies to combat the spread of viral infections.

**Significance Statement:** In the ongoing COVID-19 pandemic, we struggle to find effective control policies that “stop the spread”. While current animal models of virus spread in populations are highly sophisticated, they rarely explore effects of host behavior and its environment. We developed an experimental animal model system that allows us to visualize virus transmission in whole populations of *C. elegans* while also measuring behaviors. We were able to demonstrate how *C. elegans* genetics influences the progression of viral infection in a population and how animals adjust their behavior when infected. In the future, we envision that animal model systems like ours are used to test the effects of viral control policies on viral spread before they are applied in real world scenarios.

## Main Text

### Introduction

Throughout history, humankind has faced threats from infectious diseases, and human populations have been repeatedly shaped by pandemics. *Yersinia pestis* (the “black death”) wiped out 30-60% of Europe’s population in the 14th century, and more recently, 3-5% of the world’s population died as the result of influenza virus infections in 1918 (1). Today, the spread of viral and bacterial pathogens is ever more amplified due to the world’s unprecedented interconnectedness as seen in the recent COVID-19 outbreak (2–5). Hence, it becomes ever more important to predict and control such deadly outbreaks.

Epidemics are driven by the transmission of a pathogen from infected hosts or carriers to susceptible ones, leading to the pathogen’s spread in the population. The dynamics of these transmissions are often difficult to predict, as they result from complex and interdependent interactions between biological processes in both the pathogen and the host, the environment, and the behavior of the host population (6–13). Therefore, experimental animal models are frequently used to simulate pathogen dynamics and to facilitate the design of control policies. Laboratory mouse populations, for example, have been previously used for transmission experiments of a bacterial pathogen to directly measure the bacterial reproductive ratio through blood and fecal sampling (14, 15). In other studies, infections that lead to host death were used to measure infection levels via the host mortality rate (16, 17). A more direct and less invasive visualization of infection can be achieved by using transparent hosts. For example, microparasite infections in transparent frog embryos and fungal pathogens in *Daphnia dentifera* allowed pathogen detection *in vivo* by eye (18, 19). While these models have greatly advanced our understanding of infectious diseases, they also have some limitations: viral infections are difficult to follow over time in non-transparent hosts, and genetic tools for organisms like *Daphnia* are often limited.

This manuscript introduces the infection of the nematode *Caenorhabditis elegans* by the Orsay virus (OrV) scored by a fluorescent reporter as a novel host-pathogen model (“epidemiology-in-a-dish”). *C. elegans* is small (∼1 mm), feeds on bacteria such as *Escherichia coli*, and can be easily maintained on agar plates, thus allowing easy long-term imaging and tracking of individuals.

Discovered in immunocompromised wild *C. elegans* isolates in France in 2011, the OrV is the only natural viral pathogen of *C. elegans* found so far (20) (see review (21)). This positive-stranded RNA virus primarily infects intestinal cells (22). The infection is not lethal and results in a mild phenotype, inducing fusion of intestinal cells and slowed progeny production (20). In populations, the virus spreads only by horizontal transmissions, in a similar way to feces-to-oral transmissions: viral particles are released from the intestine of infected animals into their environment through the rectum and enter other individuals while they are feeding on the contaminated bacterial layer (20). *Wild-type* animals (N2) show no symptoms of infection, as they have two alternative pathways for viral defense, analogous to an innate and an acquired immune response. While N2 animals are asymptomatic and only carry a low viral load, some immunodeficient mutants have been identified to be susceptible to the virus. One family of mutants is deficient in the antiviral RNA interference (RNAi) response (20, 23), which relies on factors of the classical RNAi pathways (24), but also specifically requires the Dicer-related helicase (DRH)-1 protein for recognition of the viral RNAs (23). A second family is mutated in the co-suppression defective (CDE)-1 protein, which uridylates viral RNAs, a signal that leads to their degradation (25). Finally, general transcriptional responses have also been characterized that, upon infection, regulate expression of innate immunity genes (23, 26).

Our fluorescent reporter enabled us to monitor individual transmission events in real-time, allowing for subsequent quantification of both infection events and host behavior. We used a customized epifluorescence microscope setup to monitor infection dynamics in *C. elegans* populations of different genotypes and demonstrated how different genotypes have specific OrV transmission dynamics. Finally, we visualized how viral infection impacts *C. elegans* behavior. This way, the infection of *C. elegans* with the Orsay Virus enabled us to measure, manipulate and control the effects of key epidemiological parameters on infection dynamics and on host behavior with unprecedented precision.

## Results

We previously developed a fluorescent infection reporter which enables a visual read-out of OrV infection at the individual level (25). The reporter comprises two parts. To detect and track individual worms, a fluorescent protein (mcherry) is constitutively expressed by a tissue-specific promoter in the nematode pharynx. A second fluorescent protein (GFP) is under the control of an immune-response-activated promoter (*lys-3*), leading to an infection-specific fluorescent signal in the intestine. (**Fig. 1A, D, E**). We validated the reporter by comparing the fraction of GFP positive nematodes to real time PCR measurements of viral RNA, collected in the same animals (**Suppl. Fig. 1**).

**Figure 1:**
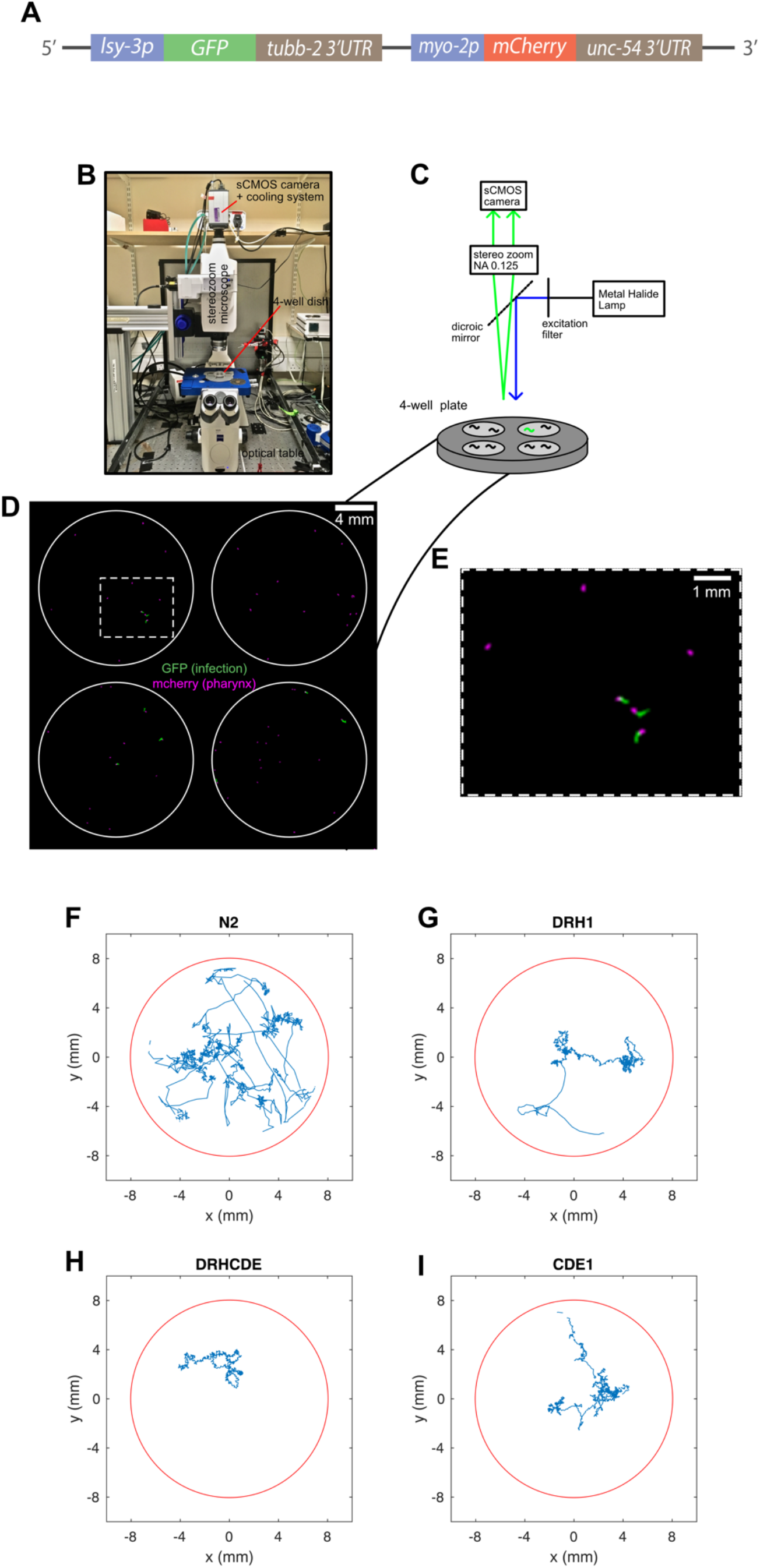
The Orsay Virus infection reporter and custom microscope setup. **(A)** Schematic representation of the infection reporter. The infection reporter used in this study consists of the [*lys-3p::GFP::tubb-2 3′UTR + myo-2p::mCherry::unc-54 3′UTR*] transgene (25). In this transgene, the GFP gene is under the control of the *lys-3* promoter that is known to be upregulated upon infection. This leads to the expression of a GFP signal in the intestine of infected individuals. The *mcherry* gene is constitutively expressed by the pharynx-specific *myo-2* promoter, leading to a permanent red signal that can be used to spot and track each individual of the population. **(B)** Picture of the microscope setup with a mounted 4-well plate. **(C)** Cartoon depiction of the customized microscope setup. In each experiment a single plate with 4 agar filled wells is imaged for 2-3 days with frame rates of 10-20 sec to score both infection and location of individual nematodes. **(D)** Snapshot of a post-processed fluorescence image of the 4-well plate containing ∼20 worms per well. Wells are highlighted by white circles and worm pharynx are shown in magenta while infected worms are green. **(E)** Magnification of the highlighted area in D. **(F)-(I)** Example trajectories of nematodes for genotypes N2, cde-1, drh-1, drh-1;cde-1. The red circle represents the outline of the well the worm is imprisoned in.

Our experimental setup utilized a customized upright fluorescence stereomicroscope that can be controlled by a laptop and to allow simultaneous imaging of four spatially separated nematode populations in a 4-well plate (filled with agarose) (**Fig. 1B-C**). To highlight the influence of host genetics – in particular its susceptibility to the virus – on the dynamics of infection in populations, we worked with two characterized mutations that result in immunocompromised animals: *cde-1* (compromised in antiviral uridylation response) and *drh-1* (compromised in antiviral RNAi response). We also investigated a double mutant that carries both mutations (*drh-1;cde-1*).

Viral filtrate was homogeneously mixed with a bacterial food layer (*Escherichia coli* HB101) before seeding as done previously (27). Hence, nematodes were infected from a homogeneous environment, and individuals were expected to rapidly ingest the virus while feeding. We seeded each 16mm wide well of the 4-well plates with an HB101-OrV mixture, and an isogenic population of 20-25 L4-stage nematodes of one of the four genetic backgrounds (*wild-type (*N2*), drh-1, cde-1*, and *drh-1;cde-1*). Nematode infection was subsequently recorded over 2-3 days with images (GFP/viral load & mcherry/pharynx channels) taken every 10-20 seconds. The resulting movies were post-processed to enhance the fluorescence signal (see **Fig. 1D-E** for an example of a post-processed fluorescence image) and subsequently analyzed automatically with a custom-written script to extract animal locations, infection levels, and individual tracks (see **Fig. 1F-I**).

Our setup enabled us to follow and quantify the infection of individuals over the course of several days at high temporal resolution (**Fig. 2**). Animals were tracked over 55 hours with simultaneous scoring of infection reporter expression. We quantified infection levels by dividing the number of infected animals by the total number of animals in each frame. Different genetic backgrounds exhibited characteristic infection dynamics that were similar between two separate biological repeats (**Fig. 2B, Table 1**). OrV infection onset in both cases occurred earlier in the *cde-1* mutant than in the *drh-1* mutant and the infection spread rapidly and early on, reached a peak and then attenuated (**Table 1**). In contrast, in the *drh-1* population, the infection spread more slowly and remained active until the end of our imaging period (55h) while the *cde-1* population was able to clear the virus (**Table 1**). The double *cde-1;drh-1* mutant exhibited an intermediate behavior with similar early dynamics as the *cde-1* mutant (rapid onset of infection) and with comparable late dynamics to the *drh-1* mutant (longer persistence) (**Table 1**). In the wild-type population, some worms were GFP positive, i.e. infected, for short periods of time but there was no widespread epidemic (**Fig. 2B**).

**Figure 2:**
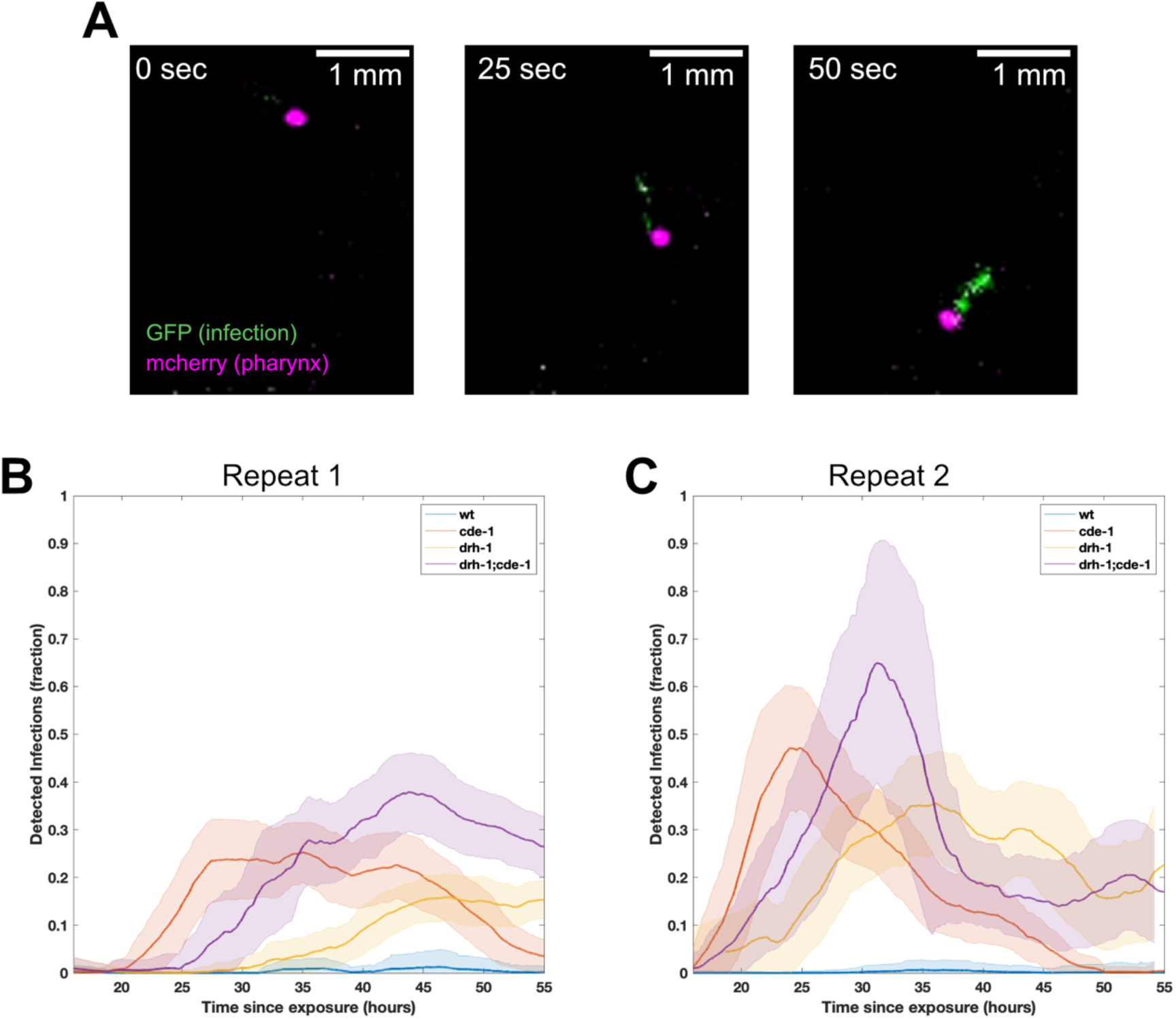
Dynamics of transmission. **(A)** Example trajectory of a nematode that becomes infected over time. The gfp signal is barely visible at t=0 sec and gradually increases over time until the maximum infection is detected at t= 50 sec. **(B-C)** Quantification of infection dynamics in WT, *cde-1, drh-1*, and *cde-1;drh-1* populations for two separate biological repeats with the same genotypes. Y-axis represents the fraction of infected worms, i.e., GFP positive ones, on the plate and X-axis the time passed since plating. The plots depict the moving average +/- standard deviation (bin size 2.8 hours). Quantifications shown in Table 1.

**Table 1:** Quantification of Infection dynamics in Figure 2B-C.

| *Experiment* | *Strain* | *Maximum*<br>*Infection (frac)* | *Initial Rate*<br>*(frac/min) over*<br>*5.5 hours from*<br>*onset* | *Recovery*<br>*(Frac of*<br>*Maximum)* | *Onset Time*<br>*(10%*<br>*infection)* |
| --- | --- | --- | --- | --- | --- |
| 1 | 'wt' | 0.01 | - | - | - |
| 1 | 'cde-1' | 0.25 | 2.00 | 0.08 | 23.9 |
| 1 | 'drh-1' | 0.17 | 0.36 | 0.99 | 40.56 |
| 1 | 'drh-1;cde-1' | 0.38 | 1.45 | 0.53 | 29.63 |
| 2 | 'wt' | 0.00 | - | - | - |
| 2 | 'cde-1' | 0.47 | 2.73 | 0.00 | 18.30 |
| 2 | 'drh-1' | 0.36 | 0.60 | 0.43 | 24.24 |
| 2 | 'drh-1;cde-1' | 0.65 | 1.32 | 0.21 | 20.25 |

Next we tested whether nematode behavior was affected by the Orsay virus by analyzing behavior pre-and post-infection for the four different genotypes, *wild-type, drh-1, cde-1*, and *drh-1;cde-1* (**Fig. 3**). Infected and uninfected worms exhibited similar behavioral changes upon infection, e.g. speeds and inter worm distances, across different genotypes (**Fig. S2**) allowing us to pool the results of the four genotypes together into one group of infected and one group of uninfected nematodes. Velocities of uninfected nematodes were significantly (quantified as effect size 95% confidence interval) higher than those of infected worms (**Fig. 3A-B**). Infected nematodes were by an average of 330 [300-365] µm/sec slower than uninfected ones in biological repeat 1 and 115 [92-138] (µm/sec) slower for biological repeat 2 (**Fig. 3A-B**). [..] signifies the 95% confidence interval of the average effect size value. The average inter-animal distance is an approximation of the population density and the “contact network” (i.e. the number of average contacts a worm has). The average inter-animal spacing was 1.43 [1.28-1.57] mm (mean effect size [95% confidence interval]) larger for infected worms compared to uninfected ones for biological repeat 1 and 1.26 [1.08-1.44] mm larger for biological repeat 2 (**Fig. 3C-D**), suggesting that uninfected worms avoid infected worms. We investigated this further by transferring uninfected worms on virus plates on which worms had been infected previously. Transferred uninfected worms avoided the food suggesting that infected worms secret an avoidance signal that is sensed by other worms (**Fig. S4**).

**Figure 3:**
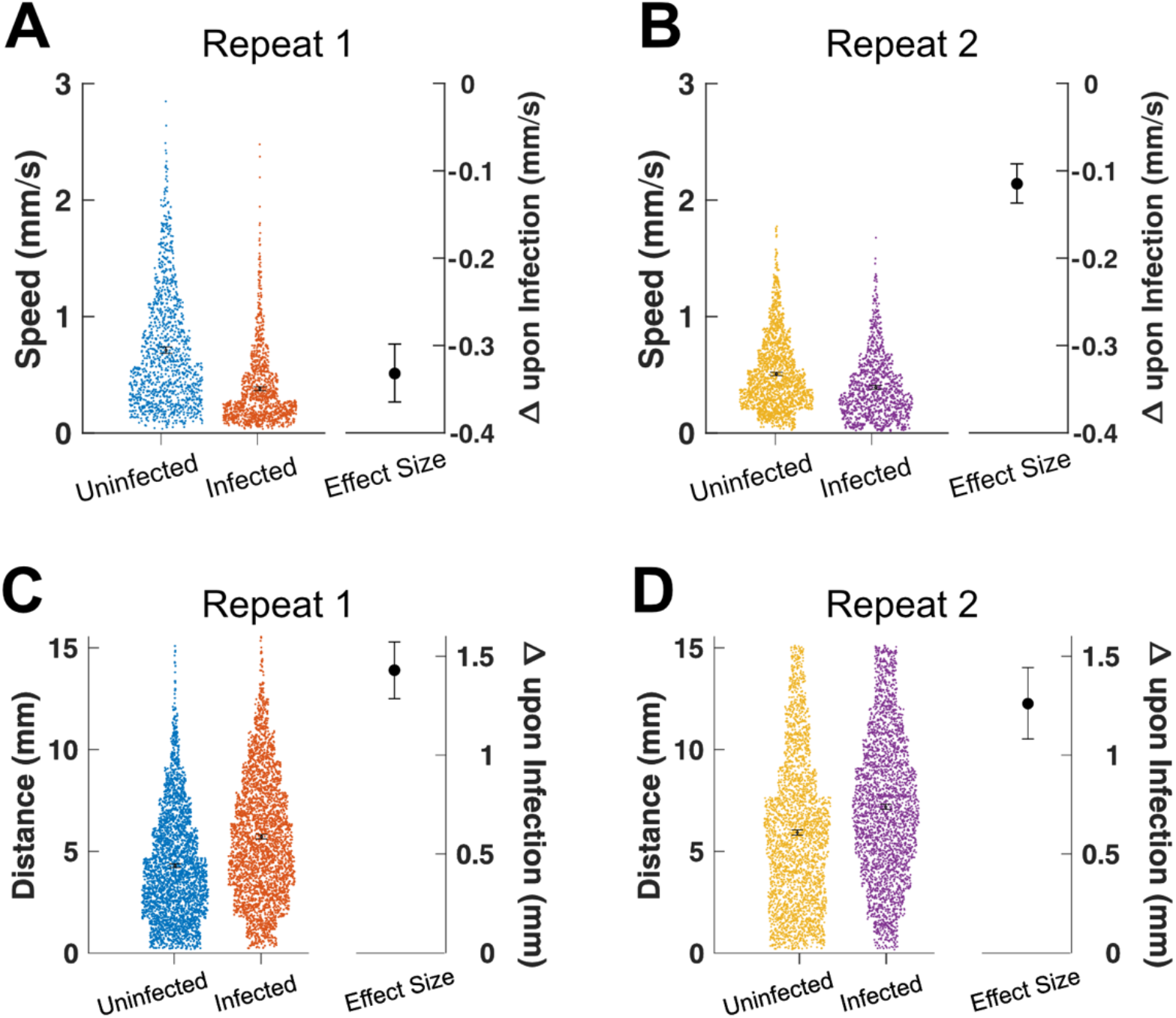
Animal behavior is changed upon infection. **(A)** Measured nematode velocities for uninfected immunocompromised worms and infected ones. Data was pooled over all strains as infected vs uninfected worm behavior was similar between genotypes (**Fig. S2**). Each dot represents one frame to frame velocity measurement. Points and intervals represent the mean and 95% confidence interval of the distribution and the effect size. **(B)** Change in nematode velocities in a separate biological repeat. **(C)** Nematode Inter-animal distance with effect size. **(D)** Inter animal distance for another biological repeat.

Our model system can be used to explore how behavior affects viral spread. Both biological repeats exhibited different infection dynamics with the onset and severity of population infection being higher in the 2^nd^ repeat (**Table 1**). Uninfected immunocompromised worms exhibited lower velocities in repeat 2 compared to repeat 1 (**Fig. S3**) suggesting that worm mobility is linked to infection dynamics.

## Discussion

Here we introduced a new “epidemiology in a dish” model to quantify viral infection spread in *C. elegans* populations and started connecting information about the behavior of individual uninfected and infected worms with the dynamics of infection of the whole population. In the future, parameters that potentially contribute to viral spread, such as nematode behavior, environment, and genetic heterogeneity, can be selectively manipulated and their effect on viral transmission dynamics be tested.

We observed that *cde-1* mutants exhibited an earlier onset of infection than *drh-1* mutants (**Fig. 2B-C, Table 1**). This suggests that the antiviral uridylation response, which is downregulated in *cde-1* mutants, provides an earlier immunity response (also supported by (25)), while *drh-1* mediated antiviral RNAi is important for later stages of infection. We speculate that the RNAi response is delayed due to effectors, i.e. the small viral RNAs, that need to be synthesized before DRH-1 can recognize them. The double mutant (*drh1;cde-1*) exhibited an early onset of infection combined with a longer persistence of infection than the *cde-1* mutant background, indicating that *cde-1* and *drh-1* are indeed part of independent pathways (25).

In addition to host immunity and genetic background, host behavior is also known to influence the dynamics of infectious disease transmission. Host mobility and relative population density, in particular, shape the population’s contact network, i.e. the frequency of contact between individuals, which the infection needs in order to spread. The role of direct contacts in infection spread of the OrV is currently unknown, but co-localization of individuals may enhance transmission even without direct contact by increasing the likelihood of individuals coming into contact with shed virus. Nematodes slowed down upon infection while inter-animal distances increased (**Fig. 3A**). Both effects combined are likely to decrease the rate at which infected and uninfected individuals co-localize, potentially decreasing viral spread. We speculate that the behavioral response is intended to mitigate viral spread, as we have observed that uninfected worms can exhibit avoidance behavior towards food patches exposed to infected animals (**Fig. S3**). Similar observations have been made in other organisms, where infection induced a characteristic “sickness behavior” (28). Our current algorithm is able to reliably track worms over an average of 1 hour.

We observed that infection progressed differently in populations with different behaviors (**Fig. S3**). In future work, nematode behavior, such as motility and worm aggregation (the frequency of contact between individuals), can be directly perturbed to measure its effects on viral spread. Worms can be slowed down, for example, by using strains with abnormal locomotion phenotypes, such as *unc* (29) or *rol* (30) mutants, as well as mutants for neurotransmitters that are known to affect locomotion rate by rendering worms unable to sense food or their movement more erratic (31). In particular, dopamine-defective mutants (32) could be used to study the observed slowing down upon infection, as the dopaminergic pathway is thought to induce the reduction in locomotion characteristic of this response (33). In addition, other mutants of *C. elegans*, such as *npr-1* (a mutant with an aggregation phenotype (34)), could be used to directly change the frequency and the group size of nematode social interactions. This would serve as an approach to measure the effects of social interactions on viral spread. Additionally, fluorescence reporters could be incorporated into the virus itself, to allow for direct visualization of both the virus in the organism and of viral concentrations outside of the animal including virus shed into the food source.

We here visualized viral spread in a population whilst quantifying host behavior. The latter plays an equally important role as the underlying immune response as became apparent in the ongoing COVID-19 pandemic in which mitigation focused on behavior. We envision our system as a complementary experimental model to probe how behavior and immunity integrate to combat viral transmission potentially informing new policies to combat global pandemics.

## Materials and Methods

### Genotypes

Strain N2: wild-type genotype from Brenner - CGC

Strain JU1580: wild isolate in which the OrV has been discovered. It carries a polymorphism of the *drh-1* gene (Isolated by Marie-Anne Félix (20) – Imported in the Miska lab by Alyson Ashe).

Strain SX2635 *mjIs228 ?*. This strain carries the infection reporter in a N2 background. The question mark indicates that the position of the allele in the genome is currently unknown. Source: Miska lab (Jérémie Le Pen) (25).

Strain SX2790 *drh-1(ok3495)* IV; *mjIs228 ?*. This strain carries the infection reporter and the *drh-1* mutation. Source: Miska lab (Jérémie Le Pen) (25).

Strain SX2999 *cde-1(tm1021)* III; *mjIs228 ?*. This strain carries the infection reporter and the *cde-1* mutation. Source: Miska lab (Emma Kneuss) (25).

Strain SX3017 *cde-1(tm1021)* III; *drh-1(ok3495)* IV; *mjIs228 ?*. This strain carries the infection reporter and both the *cde-1* and the *drh-1* mutation. Source: Miska lab (Emma Kneuss) (25).

### Preparation of viral filtrates

A stock of viral filtrates was prepared using a liquid culture protocol from David Wang’s lab.

Firstly, “thick” bacteria food was prepared as follows. HB101 bacteria were inoculated in 30 ml of SuperBroth (1.3% Bacto-tryptone, 2.6% Yeast Extract, 0.4% glycerol) supplemented by K-orthophosphates (0.17 M KH2PO4, 0.72 M K2HPO4) (SuperBroth-K) and shaken at 200 rpm for several hours at 37°C. 5 ml of this starter culture were then transferred to 1 l of SuperBroth-K and shaken at 200 rpm overnight at 37°C. Next, the cultures were centrifuged at 4000 rpm for 15 min at 4°C (Sorvall RC 3B Plus centrifuge, rotor H-6000). The supernatant was then discarded, and the pellets were re-suspended in 30 ml of sterile H_2_O.

In parallel, three young JU1580 adults were transferred on 50 mm NGM agar plates seeded with HB101. 20 µl of a previous viral filtrate were pipetted onto the bacterial layer in drops and the worms were incubated at 20°C for 3 to 4 days, to let them multiply. Just before getting starved, worms were chunked onto 90 mm plates and incubated at 20°C. Just before starvation, the worms were washed off the plates with 5 ml of S-medium (prepared as described in (35)) and transferred to 250 ml of S-medium previously mixed with 30 ml of thick bacteria food. These liquid cultures were then incubated at 20°C and shaken at 160 rpm for 6 days (the cultures were supplemented with 10 ml of thick bacteria food after 3 days). To collect the supernatant, the cultures were let on ice for 60 min for the worms to settle down. The supernatant of the cultures was then collected and centrifuged at 16,000g for 30 min (J6-MI Centrifuge – Beckman Coulter). Finally, the supernatant was cleaned by filtration on a 0.45 µm pore-size membrane then on a 0.22 µm membrane (Millipore). The total volume was aliquoted, frozen in liquid nitrogen and stored at −80°C.

Finally, the new viral filtrate stock was tested by comparing the level of infection achieved in JU1580 worms with the one of the previous stocks. The new viral stock was as competent as the previous one (data not shown).

### Infection protocol

The Orsay virus filtrate was diluted within the HB101 bacterial food (1:10) to ensure homogeneous distribution of the virus on the plate. The mixture was then seeded on NGM agar plates. These plates, referred to as “virus plates”, were left to dry at room temperature for 2-3 days and then stored at 4°C for 1 month maximum. Before use, plates were kept at room temperature for at least one hour. In parallel, mock plates were prepared by diluting M9 medium into the HB101 bacterial food (1:10).

### RT-qPCR and viral load measurements

Worms were harvested using M9 medium supplemented with 1:100 worm lysis buffer (to avoid worms to stick within the tip) and let on ice to settle down. Supernatant was removed and the worm pellet was washed 3 times with M9 medium only. After the washes, most of the M9 medium was removed and the pellets were frozen in liquid nitrogen and stored at −80°C. The lysis was performed in PCR strips by transferring 5 µl of worm pellet into 45 µl of Lysis Solution supplemented with 1:100 DNase I (Ambion). The samples were then lysed by at least 10 freeze-thaw cycles using liquid nitrogen and a warm water bath, followed by 30 min of vortexing at room temperature. Lysis was stopped by adding 5 µl of Stop Solution (Ambion). Retro-transcription (RT) and qPCR reactions were then performed using the Power SYBR Green Cells-to-Ct kit (Ambion), according to the manufacturer’s instructions. Either 5 or 10 µl of worm pellet were used in the RT reaction (depending on the pellet size) and the cDNA samples were stored at −80°C. 2 µl of a 1:10 dilution of the cDNA were used as template in the qPCR reactions, which were run on a StepOne Plus qPCR machine (Applied Biosystems) following this program: 95°C for 10 min, 40 cycles of 95°C for 15 sec and 60°C for 1 min. Orsay virus RNA1 was amplified using the primers GW194 and GW195 (20). Samples were normalized to *gapdh* mRNA, using primers developed by (36). Primers sequences are provided in Table 3.2.

### Manual scoring

The populations were observed under a Leica M165 FC fluorescent microscope. For each replicate, the number of [RFP-positive;GFP-positive] and [RFP-positive;GFP-negative] individuals were scored. The percentage of infected worms in each replicate was then calculated as the number of GFP-positive worms divided by the total number of counts in the population.

The GFP is continually expressed in opening areas, such as the rectum, the vulva and the pharynx. As being continually exposed to environmental pathogens, these areas are thought to be under permanent immune activation. When an immunocompromised mutant is exposed to the virus, the GFP signal becomes activated in the anterior part of the intestine. On the other hand, the RFP signal is constitutively expressed under any conditions in the pharynx, in both wild-type and mutant backgrounds.

### Reliability

To assess the reliability of our reporter, we infected populations of 40 eggs of each genotype in 2 independent experiments (with n=3) for *drh-1* and *cde-1;drh-1*, and 3 independent experiments (with n=3) for wt and *cde-1*. After 3 or 4 days, we manually counted the number of GFP-positive worms in the population and, directly after, collected the population to measure its viral load by RT-qPCR on the viral genome. The RT-qPCR data are expressed as the logarithm of the fold change (ΔCT) between the Orsay virus RNA1 and the *gapdh* mRNA. Primers sequence: OrV RNA1 Forward 5’- ACCTCACAACTGCCATCTACA-3’, OrV RNA1 Reverse 5’- GACGCTTCCAAGATTGGTATTGGT-3’ (20), *gapdh* Forward 5’- TGGAGCCGACTATGTCGTTGAG-3’, *gapdh* 5’-Reverse GCAGATGGAGCAGAGATGATGAC-3’

### Long term imaging

Adult *C. elegans* from the 4 genotypes were transferred to 4 well plates with each well containing a food drop with HB101 bacteria that were mixed with OrV. 15h post plating the plates were transferred under a customized Zeiss Axio Zoom microscope which was able to take high resolution images of all 4 wells (**Fig 1D-E**). Both mcherry and GFP channels were than acquired with exposure time 1 and 5 seconds respectively every 10-20 seconds for ∼40h. Room temperature was kept at 20 degrees. After that time the next generation of larvae hatched, and the food tended to dry out.

### Image Analysis

Movies were quantified as follows:

1. The ImageJ Rolling Ball Average plugin was applied individually to both the mcherry (ball size 10 pixels) and gfp (50 pixels) channels to reduce background noise. This helped to remove large bright areas in particular.
2. Thresholding. GFP: For each frame the mean projection of the surrounding 300 frames was subtracted to reduce background variations. Subsequently, a TopHatTransformation with disk size 20 pixels was applied (built in *Mathematica* function). GFP channels were then binarized with the threshold being 7*MeanBackground. Resulting binary images were dilated with pixel size 1 and subsequently eroded with pixel size 1. MCH: Images were binarized without post processing (5.5xMeanBackground).
3. Component filtering. GFP: Only keep components that have Area>15pixel^2^ & Area<100pixel^2^. MCH: Area>7 pixel^2^& Area< pixel^2^.
4. Binarized components in the mcherry channel were linked over frames (tracked) with an adjusted version of the tracking script used in (37). This approach enabled us to track individual worms for approximately 1h. Worms were assumed to be infected if they exhibited an GFP positive signal for at least 50 frames.

### Statistics and Plotting

Plots such as those in Figure 3 and in Supplemental Figures 2 and 3 are known as effect size plots, or Gardner Altman plots (38, 39). These plots include a scatter plot of the raw data and indicate the mean of each sample. From the raw data, a collection of bootstrap samples is taken and a distribution of the effect size, in this case the difference, is estimated. The most likely effect size, and its 95% confidence interval are then calculated from this distribution. These are indicated with the black dot and the whiskers on the right of each plot. These plots were generated using the code provided (39), obtained from GitHub and run in Matlab.

## Acknowledgments

We would like to thank Zeiss for giving us access to their microscope control software.

The authors acknowledge funding by the Wellcome Trust (Research Grant 109145/Z/15/Z to M.A.H.J.), the Philippe Wiener – Maurice Anspach Foundation (postgraduate fellow scholarship to L.P.), the European Research Council (consolidator grant 772426 to K.F.), the Alexander von Humboldt-Foundation (Alexander von Humboldt Professorship to K.F.), This work was supported by Cancer Research UK (C13474/A18583, C6946/A14492) and the Wellcome Trust (104640/Z/14/Z, 092096/Z/10/Z) to E.A.M. For the purpose of Open Access, the author has applied a CC BY public copyright licence to any Author Accepted Manuscript version arising from this submission.

## Supplemental Figures

**Supplemental Figure 1:**
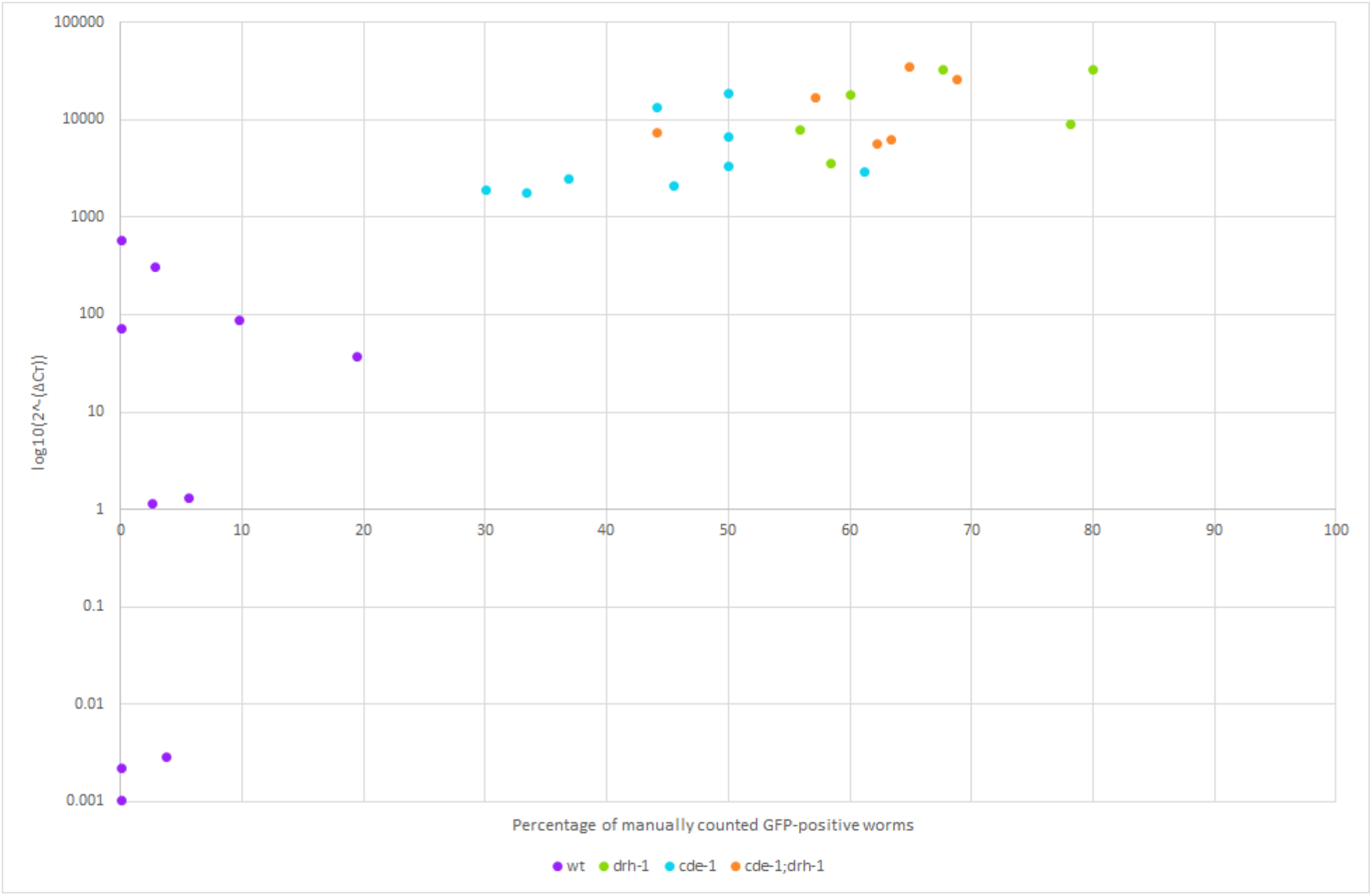
Reliability of the infection reporter. Correlation plot between the fraction of GFP-positive worms per plate (x-axis, counted by eye) and the viral load of the same population as measured by RT-qPCR (y-axis, expressed in logarithm of the fold change (ΔCT) between the Orsay virus RNA1 and the *gapdh* mRNA). Dots represent individual plates.

**Supplemental Figure 2:**
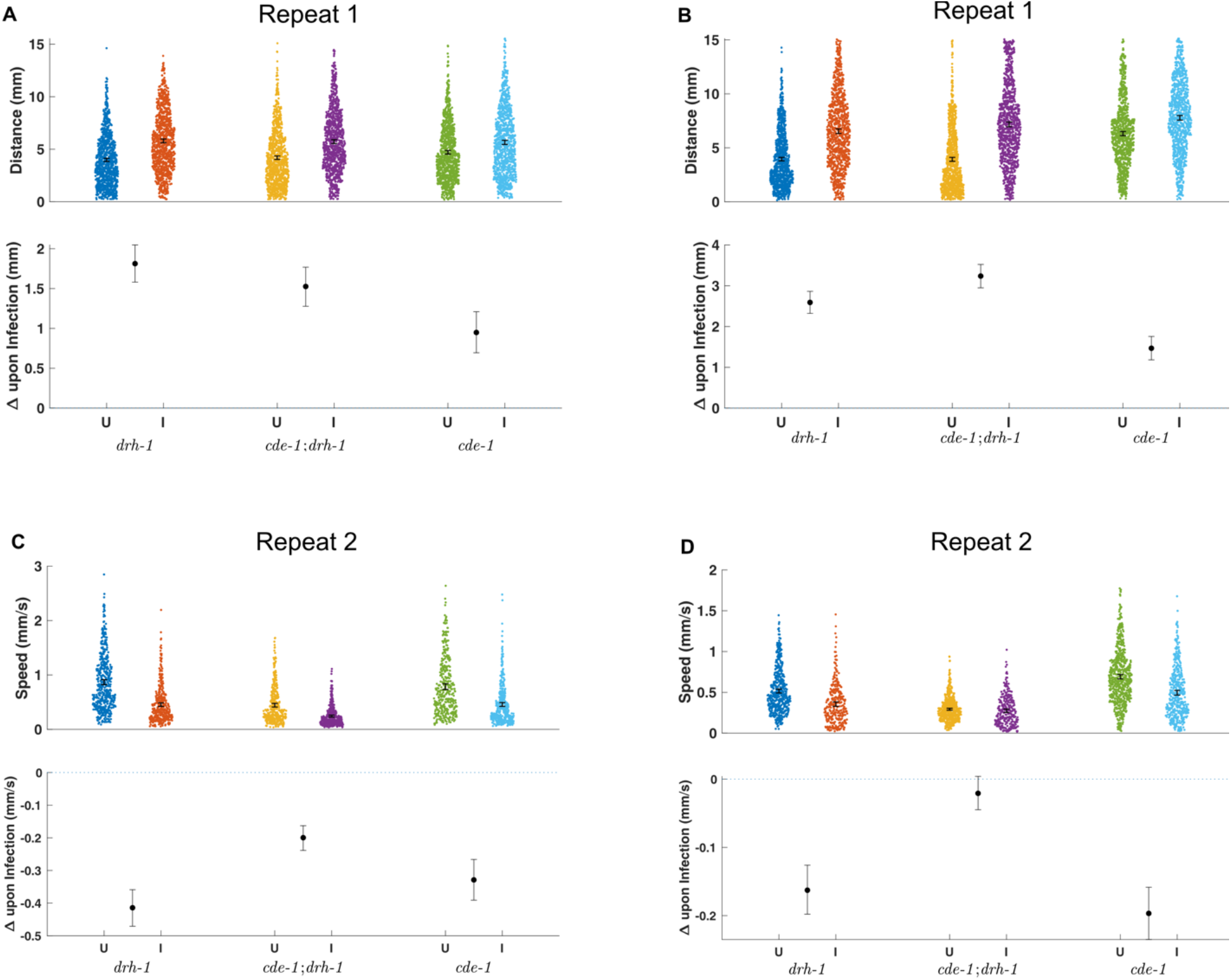
Nematode behavior changes upon infection for different genotypes in two separate repeats. **(A)** Measured nematode velocities for uninfected immunocompromised worms and infected ones. Each column represents one immunocompromised genotype and each dot represents one frame to frame velocity measurement. Points and intervals represent the mean and it’s 95% confidence interval of the distribution and the effect size. **(B)** Nematode Inter-animal distance with effect size (same as in A). **(C)** Change in nematode velocities in a separate biological repeat. **(D)** Inter animal distance for another biological repeat.

**Supplemental Figure 3:**
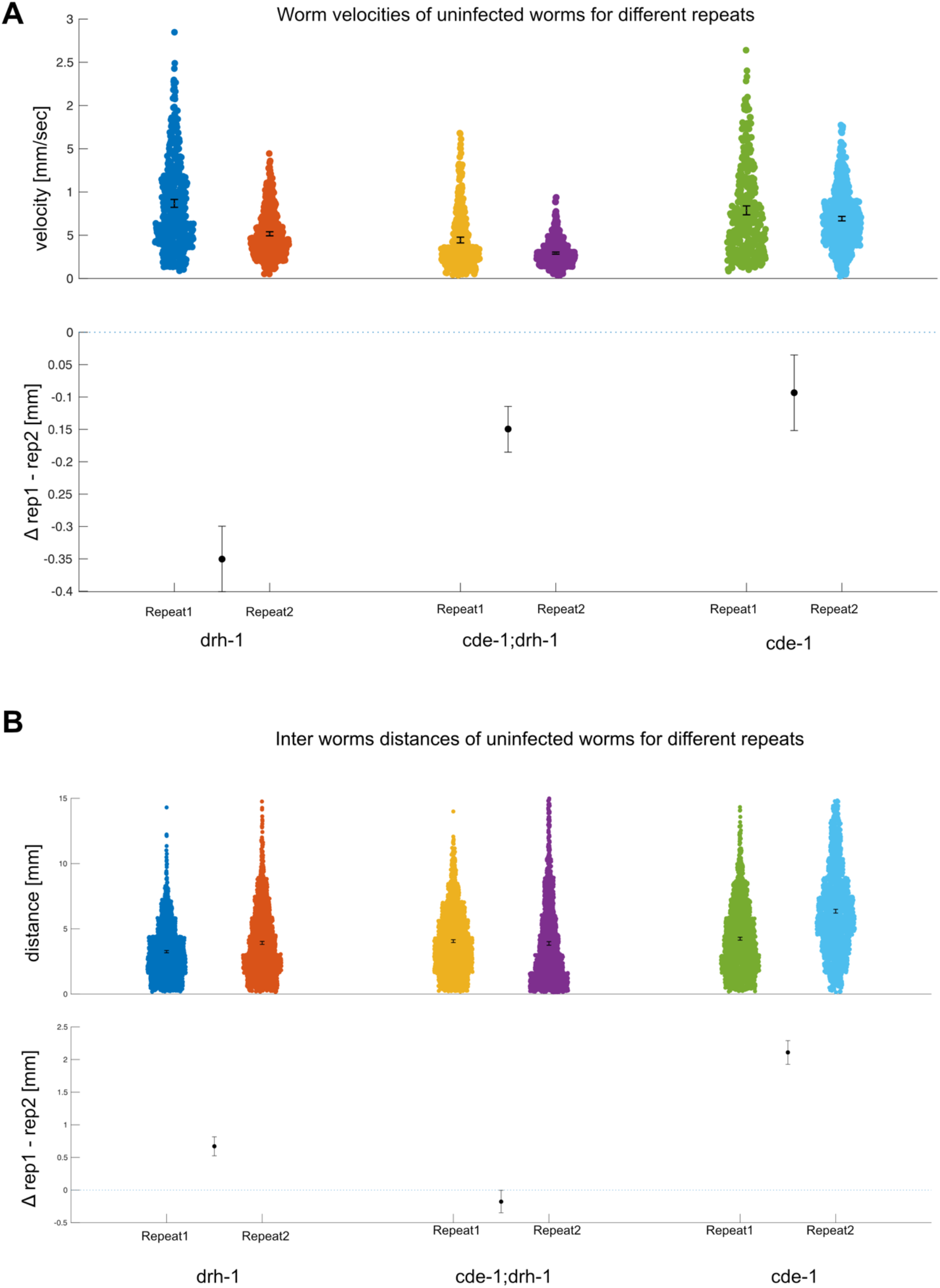
Uninfected nematode behavior compared between two biological repeats. **(A)** Measured nematode velocities for uninfected immunocompromised worms compared for repeat 1 and 2. Each column represents one immunocompromised genotype, and each dot represents one frame to frame velocity measurement. Points and intervals represent the mean and it’s 95% confidence interval of the distribution and the effect size. Bottom row represents the effect size at 95% significance. **(B)** Measured nematode to nematode distances for uninfected immunocompromised worms compared for repeat 1 and 2. Each column represents one immunocompromised genotype, and each dot represents one distance measurement. Points and intervals represent the mean and it’s 95% confidence interval of the distribution and the effect size. Bottom row represents the effect size at 95% significance.

**Supplemental Figure 4:**
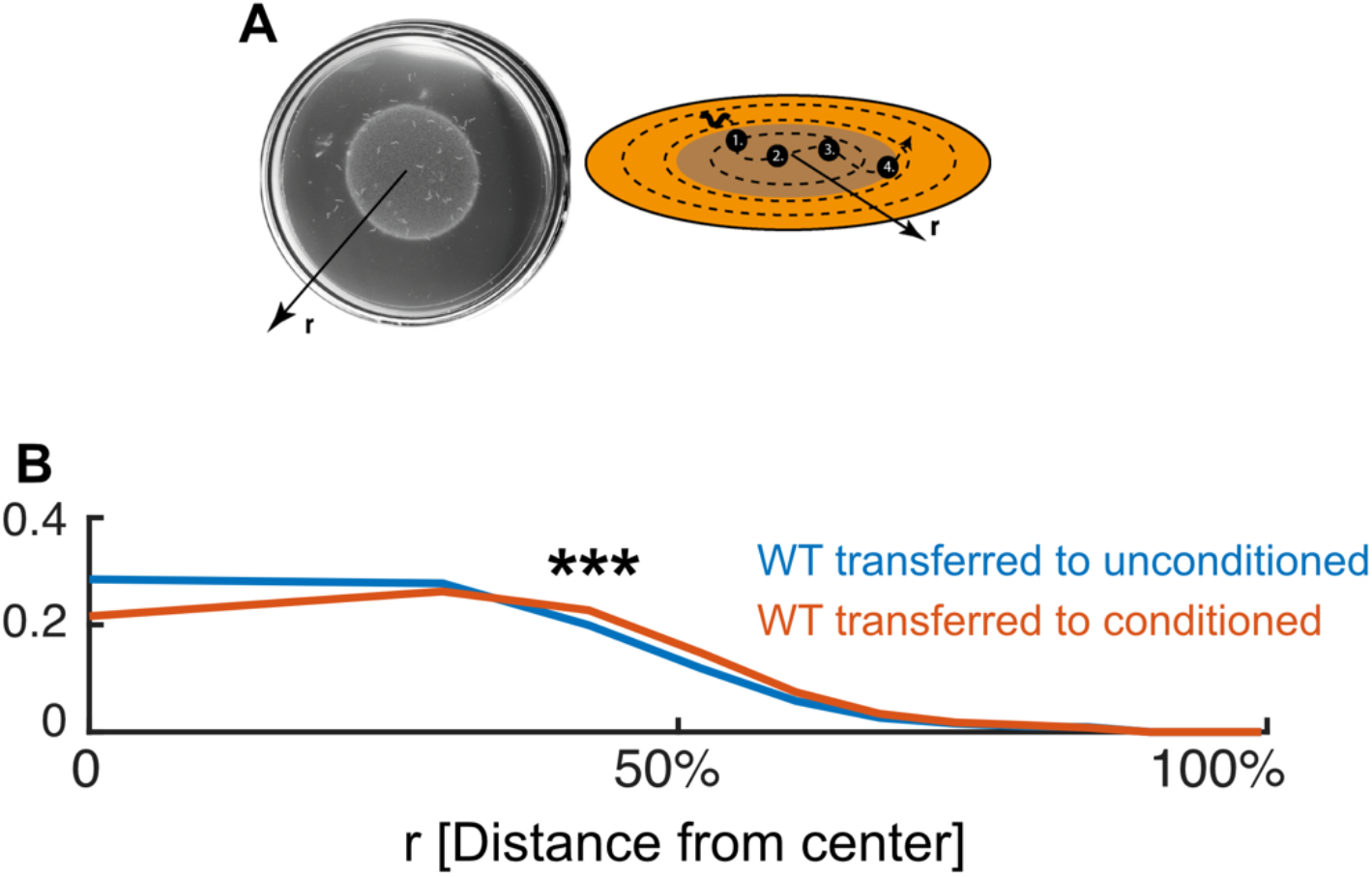
Wild type behavior on conditioned plates. **(A)** Experimental setup. Immunocompromised worms were raised on plates with either Orsay virus or normal food for 3-4 days. Subsequently the worms were removed and replaced by wild-type worms and their radial distance from the plate center measured over the course of 30 minutes. **(B)** Radial distribution of transferred wild-type worms. Distance from the dish center is given as a percentage of distance to the dish boundary. p<0.001 (Wilcoxon Ranksum Test).

## Notes

### Competing Interest Statement

The authors have declared no competing interest.

